# In silico prediction of COVID-19 test efficiency with DinoKnot

**DOI:** 10.1101/2020.09.11.292730

**Authors:** Tara Newman, Hiu Fung Kevin Chang, Hosna Jabbari

**Affiliations:** Department of Computer Science, University of Victoria, Victoria, BC Canada

## Abstract

The severe acute respiratory syndrome coronavirus 2 (SARS-CoV-2) is a novel coronavirus spreading across the world causing the disease COVID-19. The diagnosis of COVID-19 is done by quantitative reverse-transcription polymer chain reaction (qRT-PCR) testing which utilizes different primer-probe sets depending on the assay used. Using in silico analysis we aimed to determine how the secondary structure of the SARS-CoV-2 RNA genome affects the interaction between the reverse primer during qRT-PCR and how it relates to the experimental primer-probe test efficiencies. We introduce the program DinoKnot (Duplex Interaction of Nucleic acids with pseudoKnots) that follows the hierarchical folding hypothesis to predict the secondary structure of two interacting nucleic acid strands (DNA/RNA) of similar or different type. DinoKnot is the first program that utilizes stable stems in both strands as a guide to find the structure of their interaction. Using DinoKnot we predicted the interaction of the reverse primers used in four common COVID-19 qRT-PCR tests with the SARS-CoV-2 RNA genome. In addition, we predicted how 12 mutations in the primer/probe binding region may affect the primer/probe ability and subsequent SARS-CoV-2 detection. While we found all reverse primers are capable of interacting with their target area, we identified partial mismatching between the SARS-CoV-2 genome and some reverse primers. We predicted three mutations that may prevent primer binding, reducing the ability for SARS-CoV-2 detection. We believe our contributions can aid in the design of a more sensitive SARS-CoV-2 test.

**Author summary:** The current testing for the disease COVID-19 that is caused by the novel cornonavirus SARS-CoV-2 uses oligonucleotides called primers that bind to specific target regions on the SARS-CoV-2 genome to detect the virus. Our goal was to use computational tools to predict how the structure of the SARS-CoV-2 RNA genome affects the ability of the primers to bind to their target region. We introduce the program DinoKnot (Duplex interaction of nucleic acids with pseudoknots) that is able to predict the interactions between two DNA or RNA molecules. We used DinoKnot to predict the efficiency of four common COVID-19 tests, and the effect of mutations in the SARS-CoV-2 virus on ability of the COVID-19 tests in detecting those strains. We predict partial mismatching between some primers and the SARS-CoV-2 genome but that all primers are capable of interacting with their target areas. We also predict three mutations that prevent primer binding and thus SARS-CoV-2 detection. We discuss the limitations of the current COVID-19 testing and suggest the design of a more sensitive COVID-19 test that can be aided by our findings.

## Introduction

SARS-CoV-2 is a respiratory disease spreading across the world that originated in Wuhan, China in 2019 [1]. SARS-CoV-2 is most commonly detected using qRT-PCR on samples collected by nasopharyngeal swabs [2]. During qRT-PCR, the reverse primer first binds to the positive sense SARS-CoV-2 RNA genome so that the reverse transcriptase (RT) can use the primer to generate the complementary DNA (cDNA) of the negative sense strand as shown in Fig 1 [3].

**Fig 1.**
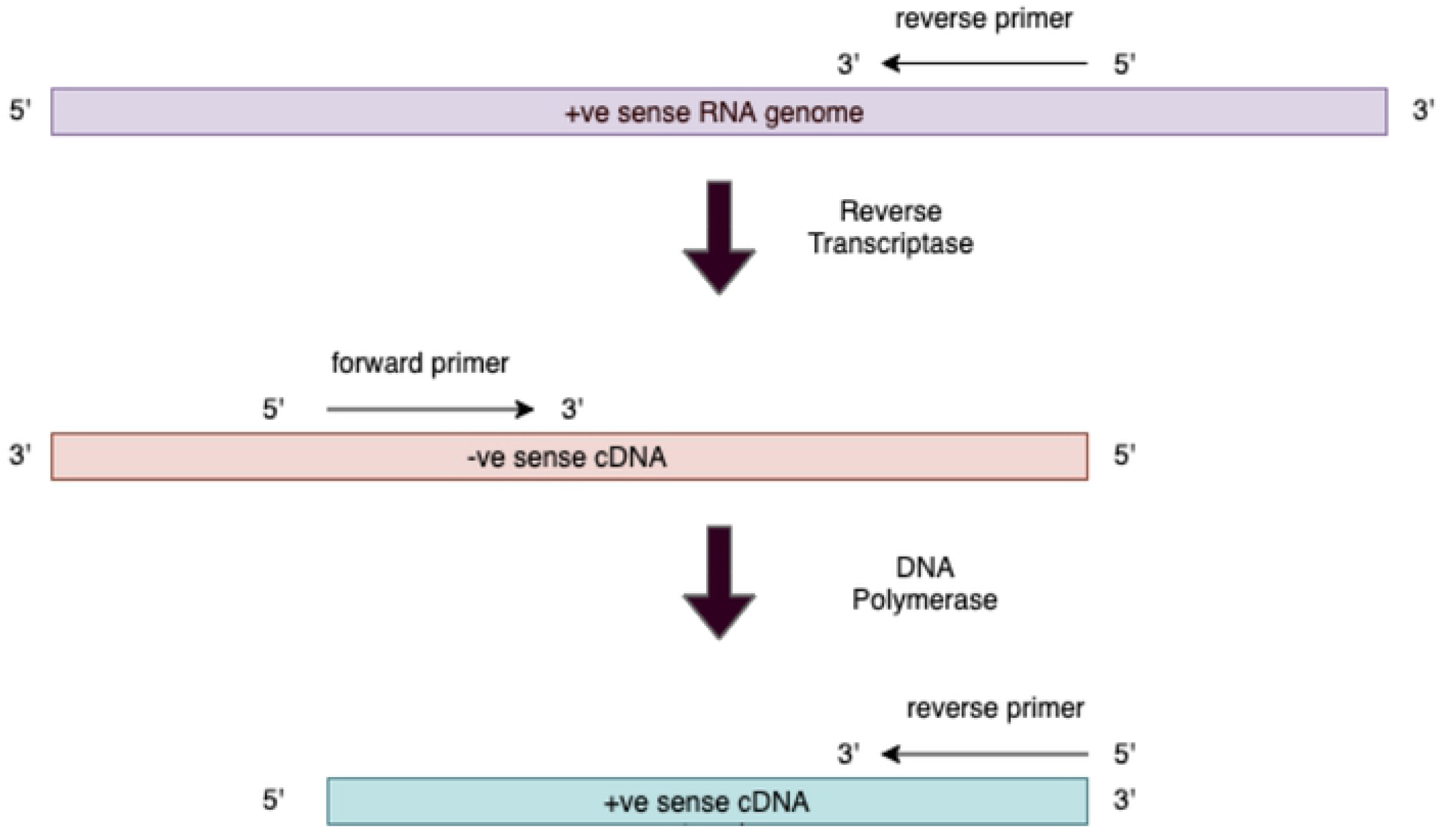
Interaction of the primers to the RNA/cDNA strands during qRT-PCR amplification. The reverse primer first binds to the target complementary sequence on the SARS-CoV-2 positive (+ve) sense RNA genome. The reverse transcriptase then generates the negative (-ve) sense complementary DNA (cDNA) strand. The forward primer then binds to the negative sense cDNA strand and the DNA polymerase generates the positive sense cDNA strand. The reverse primer binds to the complementary target sequence on the positive sense cDNA and the DNA polymerase generates a new negative sense cDNA strand. [3]. This process repeats for strand amplification during qRT-PCR.

The primer-probe set used to detect SARS-CoV-2 depends on which qRT-PCR assay is used. The China Center for Disease Control (China CDC), United States CDC (US CDC), Charité Institute of Virology, Universitatsmedizin Berlin (Charité), and Hong Kong University (HKU) have developed the most common assays to detect SARS-CoV-2 that all target different areas of the SARS-CoV-2 genome [4]. In a recent publication, Vogels *et al*. compared the analytical efficiencies and sensitivities of the primer-probe sets used in these assays [4].

By studying how the reverse primer interacts with the RNA genome, we can predict if the generation of the initial cDNA template from the RNA strand proceeds as expected during the qRT-PCR test. We do not investigate the interaction of the forward primer and the probes with the SARS-CoV-2 genome because the important interactions of these oligonucleotides are involved in binding to the cDNA, not the positive sense RNA genome.

We aimed to determine whether the RNA structure of the SARS-CoV-2 genome affects the binding of the reverse primers in the qRT-PCR assay and whether this correlated to the analytical efficiencies and sensitivities shown by Vogels *et al*. [4]. To study the structure of such interactions, we developed DinoKnot, a program that given two nucleic acid strands predicts their interaction structure. We used DinoKnot to predict how the RNA structure of SARS-CoV-2 changes when the reverse primer binds to the RNA genome and whether the primer was binding in the location that was expected. We further predicted how mutations in the primer/probe binding region of the SARS-CoV-2 genome provided by Vogels *et al*. could affect the primer/probe sensitivity for SARS-CoV-2 detection [4].

First, we introduce DinoKnot and our experiment set up. We then present the interaction structures predicted by DinoKnot. We discuss the interactions and which mutations may decrease SARS-CoV-2 detection during qRT-PCR testing. Finally, we suggest the design of a more sensitive COVID-19 test and future work that can be done using DinoKnot.

## Materials and methods

Interaction of the reverse primer and the SARS-CoV-2 genome is an example of a duplex DNA/RNA interaction, in which both strands (i.e. the RNA genome and the cDNA transcript) can be structured. Their structures can change upon interaction with one another to accommodate formation of more stable base pairings. Existing tools that predict structure of interaction in two molecules mostly focus on similar strands (i.e. DNA/DNA or RNA/RNA) [5–10], and merely focus on the interaction site (i.e. ignoring the intramolecular structures) [11–15]. Our DinoKnot (Duplex Interaction of Nucleic acids with pseudoKnots) aims to address both such shortcomings.

In this section we first provide an overview of the problem of secondary structure prediction for two interacting molecules. We then introduce DinoKnot, and explain how it overcomes the shortcomings of the existing methods on prediction of the secondary structure for two different types of nucleic acid strands. Finally we provide details of our in silico system and experiment setup.

### Secondary Structure Prediction for Interacting Molecules

An RNA is a single stranded molecule with two distinct ends, namely 5’ and 3’, that folds back onto itself by forming intramolecular base pairs. An RNA molecule is represented by a sequence, *S*, of its four bases, Adenine (A), Cytosine (C), Guanine (G) and Uracil (U) arranged on a line (representing the backbone) from 5’ (left) to 3’ (right) ends. The length of the RNA molecule is denoted by *n* and each base of the RNA sequence is referred to by its index *i*, 1 ≤ *i* ≤ *n*. Complementary bases bind and form base pairs (*A.U, C.G*, and *G.U*). Here represents a pairing of the two bases. A *secondary structure, R*, is then defined as a set of base pairs *i.j*, 1 ≤ *i* < *j* ≤ *n*; *i.j* and *k.j* can belong to the same set if and only if *i* = *k* i.e. each base may pair at most with one other base. If *i.j* and *k.l* are two base pairs of a secondary structure, *R*, such that 1 ≤ *i* < *k* < *j* < *l* ≤ *n*, then *i.j* crosses *k.l*. A *pseudoknotted* secondary structure refers to a structure with crossing base pairs. A *pseudoknot-free* secondary structure refers to a structure without crossing base pairs. The RNA structure forms because it is energetically favourable for bases to form paired helices. Different base pairing patterns in a secondary structure define different loop types. We note that the principles of structure prediction are essentially the same for single stranded DNAs. Computational secondary structure prediction from the base sequence is often done by finding the energetically most stable (minimum free energy) secondary structure, when each loop is assigned an energy value. These energy values are kept in various tables and are known as energy parameters. Some parameter sets have been derived directly from experiments, and others are extrapolated based on experimentally determined values. Energy parameters are strand type specific, i.e. similar loops in an RNA molecule have different assigned energy than the ones in a DNA molecule. Existing minimum free energy (MFE) structure prediction methods find the minimum free structure for a given sequence from the pool of all possible structures. Methods for MFE pseudoknot-free structure prediction use dynamic programming method to find the MFE structure [16, 17]. Since prediction of the MFE pseudoknotted structure is NP-hard [18, 19] and even inapproximable [20], methods for pseudoknotted MFE structure prediction focus on a restricted class of structures [21–25].

We can represent the interacting secondary structure of two nucleic acid sequences similarly, by concatenating the two strands together and keeping track of the gap between the two strands with a dummy linker. When two strands are of similar type, the energy calculation of the concatenated sequence will be similar to that of a single strand of the same type, except for loops containing the gapped region (as they are not true loops when sequences are not concatenated). When two strands of different types interact (i.e a DNA strand binding to an RNA strand) the situation is more complicated as there is currently no energy parameters available for loops formed between the two strands.

### DinoKnot

DinoKnot follows the relaxed hierarchical folding hypothesis [26] for prediction of the minimum free energy structure of two interacting nucleic acid strands. Following this hypothesis an RNA molecule first forms simple pseudoknot-free base pairs before forming more complex and possibly pseudoknotted structures [27]. During this process some of the originally formed base pairs may open up to accommodate more stable pairings. Existing methods based on hierarchical folding, namely HFold [24] and Iterative HFold [26], focus on single RNA structure prediction [24, 26]. Our DinoKnot is to the best of our knowledge the first program that follows the hierarchical folding hypothesis for prediction of pseudoknotted structure of two interacting nucleic acid molecules. Similar to HFold and Iterative HFold, DinoKnot handles a large class of pseudoknotted structures called density-2 structures [24]. This class of structures include a wide range of commonly found pseudoknotted structures including H-type pseudoknots and kissing hairpins with arbitrary nested substructures.

DinoKnot, takes a pair of nucleic acid sequences as input and returns their interaction structure with its corresponding free energy value. Each sequence can be of type RNA or DNA. We note that the minimum free energy structure of two input strands may not involve any interaction if it is energetically more favourable for each sequence to form intramolecular base pairs.

The user can optionally provide a pseudoknot-free input structure if such information is available to guide the prediction. If no input structure is provided by the user, DinoKnot will generate up to 20 pseudoknot-free secondary structures (i.e. energetically favourable stems) by default for each strand (to the total of up to 400 stems for two strands). Guided by these energetically favourable stems, DinoKnot finds multiple structures (sorted by their free energy) for the interacting structures. The output structure (in dot-bracket format) is the minimum free energy structure among this set of structures.

DinoKnot employs the energy parameters of Andronescu et al. HotKnots V2.0 [28] for RNA structures and MultiRNAFold energy parameters [6] for DNA structures. We have estimated the energy parameters for loops formed between an RNA and a DNA molecule to be the average of similar loops formed intramolecularly in an RNA and a DNA molecule.

All source code and energy parameters are available in our Github page.

### System and Experiment Setup

The SARS-CoV-2 reference genome NC_045512.2 was obtained from the National Center for Biotechnology Information GenBank database [29]. The sequence of the RNA transcripts used by Vogels *et al*. [4] to determine the primer efficiency were input into the programs Iterative HFold [26] to predict their secondary structure. This output is the secondary structure of the transcript prior to the interaction with the reverse primer. The primer sequences were obtained from the list of World Health Organization (WHO) protocols to diagnose COVID-19 [30]. The locations where the primers bind on the reference genome NC_045512.2 and the corresponding RNA transcript area are stated in Table 1.

**Table 1.**
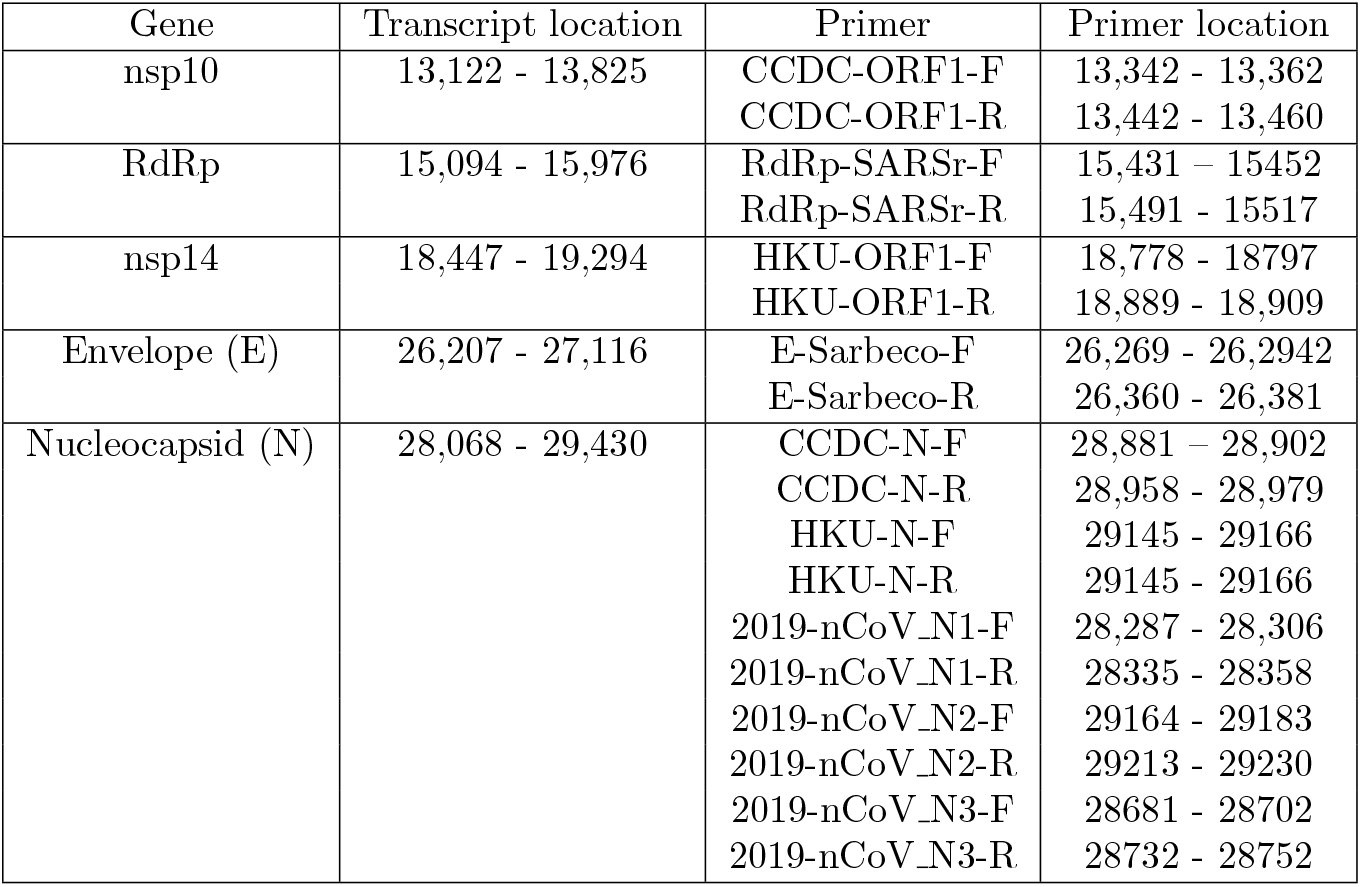
Primer binding location on the SARS-CoV-2 reference genome NC_045512.2 [29], The transcript location is the area from the reference genome input into DinoKnot to predict the reverse primer/ SARS-CoV-2 RNA genome interaction structure.

| Gene | Transcript location | Primer | Primer location |
| --- | --- | --- | --- |
| nsp10 | 13,122 - 13,825 | CCDC-ORF1-F<br>CCDC-ORF1-R | 13,342 - 13,362<br>13,442 - 13,460 |
| RdRp | 15,094 - 15,976 | RdRp-SARSr-F<br>RdRp-SARSr-R | 15,431 - 15,452<br>15,491 - 15,517 |
| nsp14 | 18,447 - 19,294 | HKU-ORF1-F<br>HKU-ORF1-R | 18,778 - 18,797<br>18,889 - 18,909 |
| Envelope (E) | 26,207 - 27,116 | E-Sarbeco-F<br>E-Sarbeco-R | 26,269 - 26,2942<br>26,360 - 26,381 |
| Nucleocapsid (N) | 28,068 - 29,430 | CCDC-N-F | 28,881 - 28,902 |
|  |  | CCDC-N-R | 28,958 - 28,979 |
|  |  | HKU-N-F | 29145 - 29166 |
|  |  | HKU-N-R | 29145 - 29166 |
|  |  | 2019-nCoV_N1-F | 28,287 - 28,306 |
|  |  | 2019-nCoV_N1-R | 28335 - 28358 |
|  |  | 2019-nCoV_N2-F | 29164 - 29183 |
|  |  | 2019-nCoV_N2-R | 29213 - 29230 |
|  |  | 2019-nCoV_N3-F | 28681 - 28702 |
|  |  | 2019-nCoV_N3-R | 28732 - 28752 |

qRT-PCR involves the use of forward and reverse primers, a probe, reverse transcriptase (RT) and DNA polymerase. Forward and reverse primers are DNA oligonucleotides that bind to complementary sequences in order to amplify a section of the SARS-CoV-2 genome. The reverse primer is a DNA oligonucleotide that interacts with both the positive sense SARS-CoV-2 RNA genome and the positive sense cDNA transcript as shown in Fig 1. Therefore the reverse primer is involved in both RNA/DNA and DNA/DNA interactions. The forward primer is a DNA oligonucleotide interacts with the negative sense cDNA transcript. The probe is a fluorescent-labelled DNA oligonucleotide that may bind to either the positive or negative sense cDNA transcript, depending on how it is designed. Therefore, the forward primer and probe are only involved in DNA/DNA interactions. RT is an enzyme that uses an RNA as a template to generate a complementary DNA strand. DNA polymerase is an enzyme that uses a DNA template to generate a complementary DNA strand.

DinoKnot was used with the RNA transcript region and the reverse primer to determine the location where the reverse primer binds and the secondary interaction structure. Two of the reverse primers, HKU-ORF1-R and RdRp-SARS-R, contain degenerate bases which means there are a mixture of oligonucleotides that contain different bases at the degenerate base position [31]. In these cases, all possible degenerate base combinations were predicted with DinoKnot. The dot-bracket output was visualized using VARNA [32] in arc format in which RNA backbone is represented by a horizontal line and base pairs are presented as arcs that connect the two bases.

#### Mutations

Vogels *et al*. listed mutations in the primer/probe binding area of the SARS-CoV-2 genome that occur at a frequency of greater than 0.1% [4]. To predict if these mutations affect primer/probe binding, and thus the sensitivity of SARS-CoV-2 detection, the mutated sequence of the transcript along with the affected primer/probe was entered into DinoKnot. The RNA/DNA setting was used to predict the interaction between the mutated RNA transcript and the reverse primer. The DNA/DNA setting in DinoKnot was used to predict the interaction structure between the mutated cDNA transcript and the corresponding primer/probe. The dot-bracket output was visualized using VARNA [32] in arc format in which RNA backbone is visualized by a horizontal line and base pairs are represented by arcs connecting bases of the backbone.

The GISAID next hCoV-19 app was used to look at the frequency of mutations in the SARS-CoV-2 genome [33].

### Data

All structure images, primer sequences and output data can be found at https://github.com/HosnaJabbari/DinoKnot_data

## Results

### Reverse primer interactions with SARS-CoV-2 genome

DinoKnot predicted the interaction structures represented in Fig 2 to occur between the reverse primer (in red) with the corresponding region around its target sequence (in green). Fig 2 represents the interaction site only. The complete structures can be found in the supplemental information on the DinoKnot Github page.

**Fig 2.**
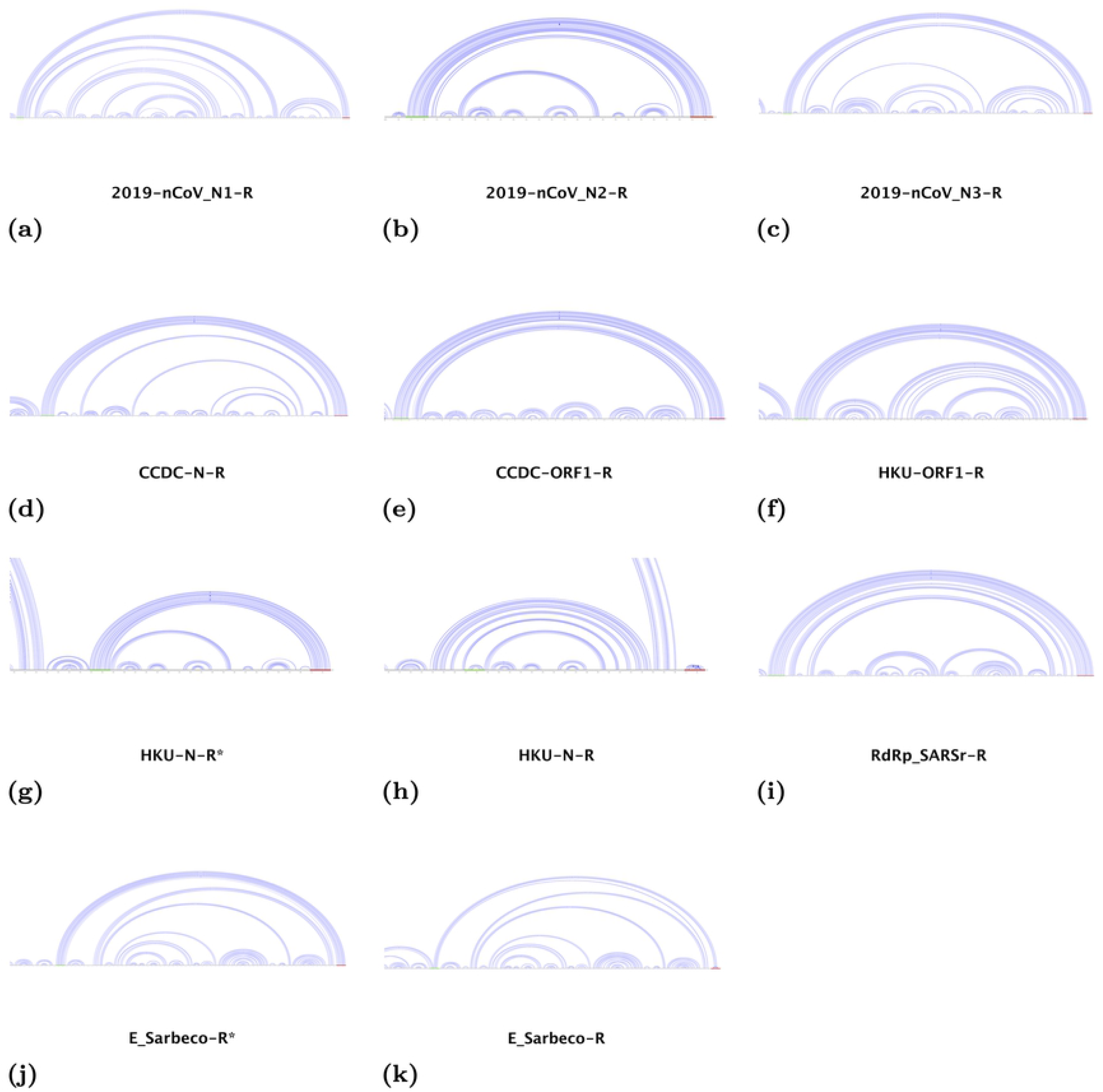
Interaction structure predicted by DinoKnot of the SARS-CoV-2 transcript genome area targeted by the reverse primer. The expected target region of the reverse primer is highlighted in green and the reverse primer sequence is highlighted in red. The nucleic acid sequences of the RNA transcript and reverse primer were input into DinoKnot to produce the output interaction structures. The E-Sarbeco-R* and HKU-N-R* primer structures were input into DinoKnot as unfolded to simulate the primer structure during the 95°C denaturation step of the qRT-PCR assay due to primer mismatch prediction under default conditions of 37° C.

**Table 2.** Energies (kcal/mol) of RNA transcript, reverse primer and reverse primer/transcript interaction structures. The transcript energy and primer energy is the energy of the transcript and primer structures before the interaction. The primer binding energy is energy of the interaction structure. the Transcript energies were predicted by Iterative HFold [26], primer energies predicted were by Simfold [5], and primer binding energies were predicted by DinoKnot.

| Primer | Transcript Energy (kcal/mol) | Primer Energy (kcal/mol) | Primer Binding Energy (kcal/mol) |
| --- | --- | --- | --- |
| 2019-nCoV_N1-R | -267.24 | -5.90 | -277.67 |
| 2019-nCoV_N2-R | -267.24 | 0 | -276.06 |
| 2019-nCoV_N3-R | -267.24 | -3.75 | -272.3 |
| CCDC-N-R | -267.24 | -1.30 | -274.67 |
| CCDC-ORF1-R | -133.39 | 0.2 | -138.54 |
| HKU-ORF1-R | -138.89 | -3.10 | -151.76 |
| HKU-N-R | -267.24 | 0* | -280.82 |
|  |  | -3.30 | -267.91 |
| RdRP-SARSr-R | -150.03 | -3.70 | -162.37 |
| E-Sarbeco-R | -148.74 | 0* | -167.39 |
|  |  | -1.50 | -150.48 |

\*E-Sarbeco and HKU-N-R primer structure forced unfolded for primer to bind to targeted area.

Vogels *et al*. experimentally found that all of the primer-probe sets had comparable analytical efficiencies that were all above 90% [4]. All primer-probe sets had comparable analytical sensitivities with a limit of detection of 100,000 SARS-CoV-2 viral copies/mL except for the RdRp-SARSr set which had the lowest sensitivity [4].

All of the reverse primers were predicted by DinoKnot to interact with their expected target region. The 2019-nCoV_N1-R, 2019-nCoV_ N3-R, CCDC-N-R and CCDC-ORF1-R primers fully paired to their target region. This prediction agrees with the analytical efficiency and sensitivity results found by Vogels *et al*. [4].

#### Partial reverse primer mismatching

The HKU-ORF1-R and the 2019-nCoV_N2-R primer were predicted to pair to their target with single base mismatches. The HKU-ORF1-R primer contains the degenerate base R which means the the primer may contain an A or a G at this position. The target base is a U in the SARS-CoV-2 genome. The interaction structure was predicted twice, both with an A and G in the position of degenerate base R. The output produced the expected match, except when the primer was set as a G, the G/U base pair did not bind but it did not affect the rest of the primer area from binding to the target area. The last base at the 3’ end of the 2019-nCoV_N2-R primer did not pair with any nucleotide but the rest of the primer was predicted to bind to its expected target region. Although these primers contain a single predicted mismatch, the analytical efficiencies and sensitivities of these primer probe sets were comparable to the other primer-probes tested experimentally [4].

#### HKU-N-R and E-Sarbeco-R

The HKU-N-R and E-Sarbeco-R primers did not pair as expected to their target region when DinoKnot was not given an input structure (i.e. when both strands were free to assume possible structures before interacting with one another). The HKU-N-R primer was predicted to bind to itself, rather than its target region. When the HKU-N-R primer structure was forced to be unfolded, the first base of the HKU-N-R primer at the 5’ end did not bind to its expected nucleotide but the rest of the primer interacted with the target region as expected. Forcing the primer to be unfolded is a prediction of the primer structure after the qRT-PCR test denaturation step at 95°C in the work done by Vogels *et al*. [4] The predicted interaction structure for the E-Sarbeco-R primer when Dinoknot was not given an input structure results in the bases at positions 8-15 in the reverse primer binding as expected to the target region. However, bases 5-7 paired with bases 16-18 and the remaining bases of the 22bp primer remained unpaired. When the primer structure was input into DinoKnot as unfolded, as done for the HKU-N-R primer, the E-Sarbeco-R primer fully paired to its target region. Both the HKU-N-R and E-Sarbeco primer-probe sets were shown experimentally by Vogels *et al*. to have analytical efficiencies and sensitivities that are comparable to the other primer-probe sets that were predicted by DinoKnot to bind completely to their target region; this supports the structures predicted with their primer structure is forced unfolded [4].

#### RdRp-SARSr-R

The RdRp-SARSr-R primer contains two degenerate bases, R and S. This means that an A or G may be present in the position of the R and a C or G may be present in the position of the S.

All combinations were given as input to DinoKnot and the predicted structures are shown in Fig 3. Fig 3 represents the interaction site only. The complete structures can be found in the supplemental information on the DinoKnot Github page. The RdRp-SARSr-R primer interacted as expected with its target region when the base at the R position was entered into DinoKnot as an A, and the base at the S position was entered as a G. The target nucleotide is a U at both of the positions where the R and S base bind. The structures were also predicted with an A input at the position of the degenerate base S. This change was suggested by Vogels *et al*. to possibly increase the sensitivity of this primer-probe set because it was found experimentally to have the lowest sensitivity of all the primer-probe sets [4]. However, this change resulted in the primer not binding to its target region.

**Fig 3.**
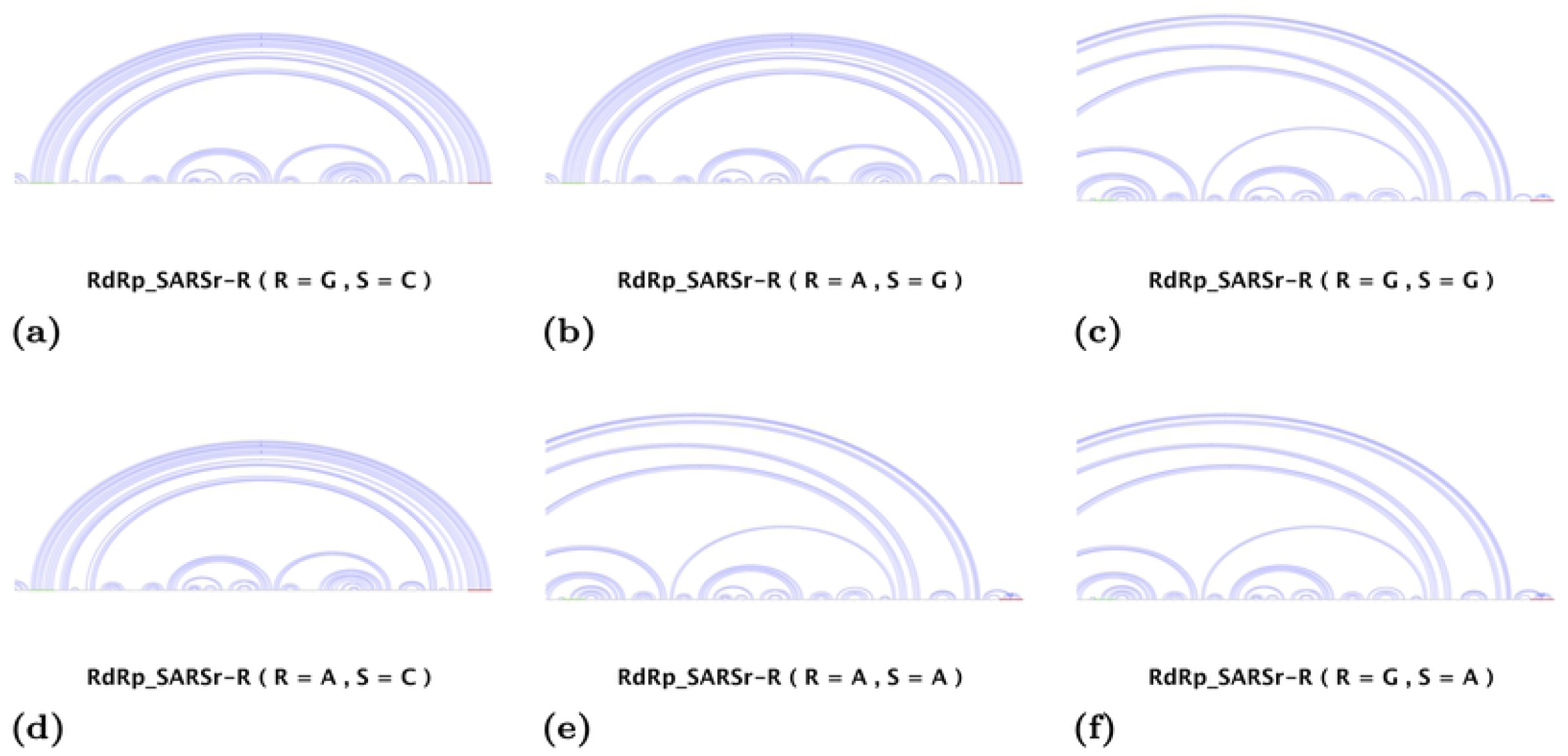
Interaction structures of the RdRp-SARS-R primer predicted by DinoKnot with all of the possible base combinations from the degenerate primers. The RdRp-SARS-R primer contains the degenerate bases R and S, which means an A or G may be present at the R position and a C or G may be present at the S position. All possible combinaions were predicted. The S position was also input as an A to predict if this change may increase primer sensitivity.

### Mutations in primer/probe binding regions

The mutations listed by Vogels *et al*. are shown in Table 3, along with the interaction structure energies [4]. The resulting structures are shown in Fig 4. Fig 4 represents the interaction site only. The complete structures can be found in the supplemental information on the DinoKnot Github page. The effects of the mutation on the interaction structure are compared to the primer binding ability of the primer/probes to the reference genome NC_045512.2. All primer/probes paired as expected to the target area of the reference genome with the exception of the 2019-nCoV_N3-F primer which has one unpaired base at the 3’ end.

**Fig 4.**
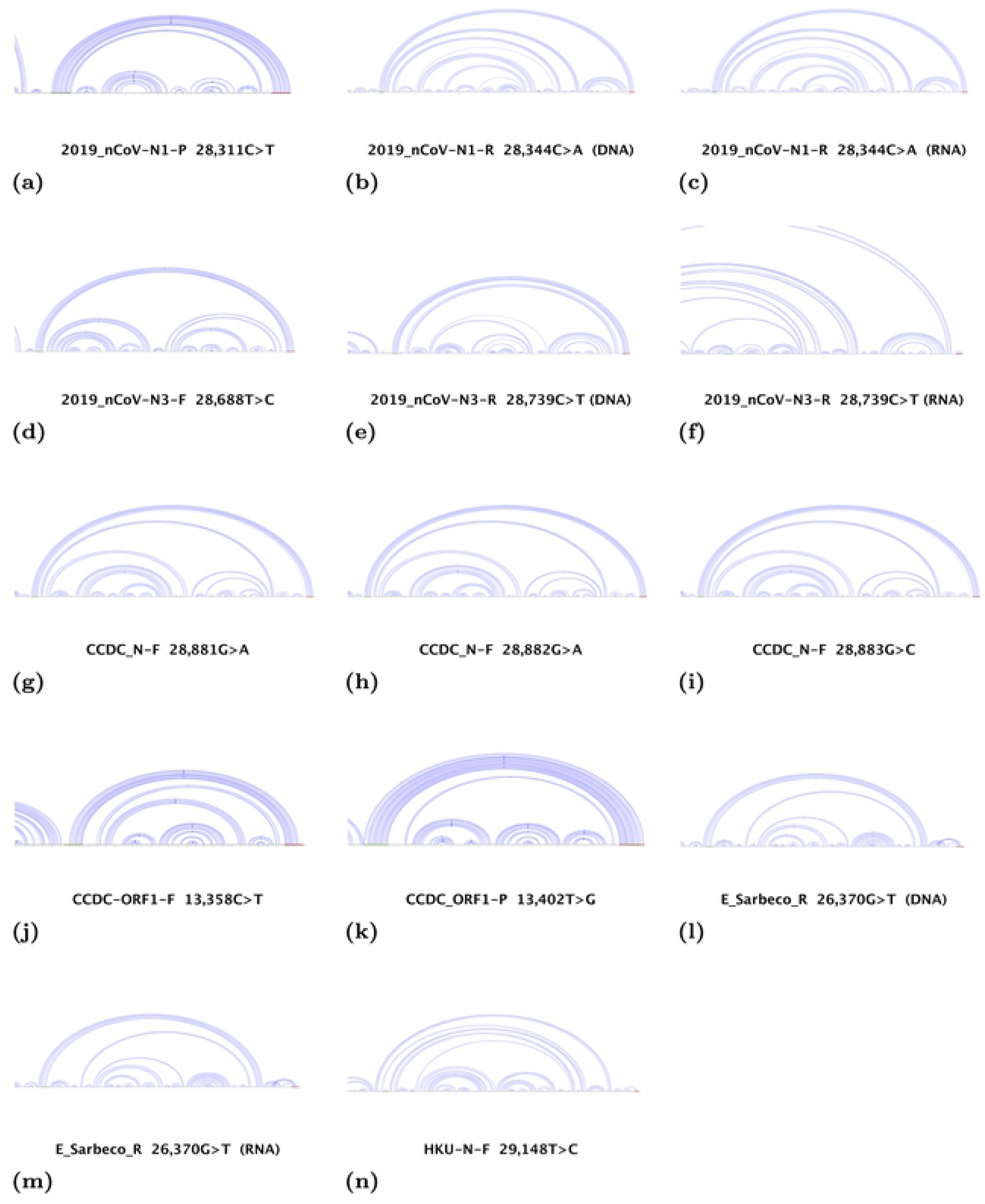
Interaction structures of primer/probes with the transcript regions containing mutations in the primer/probe binding region predicted by DinoKnot. The expected target region is highlighted in green and the primer/probe sequence is highlighted in red. The mutated base is highlighted in pink.

**Table 3.** Structure energy differences predicted with mutations in the primer binding region of the SARS-CoV-2 genome obtained from Vogels *et al*. [4]. The primer binding energy was predicted by DinoKnot. The probe/transcript and forward primer/transcript interactions are DNA/DNA interactions since these oligonucleotides interact with the cDNA strands [3]. Mutations in the reverse primer binding region include both DNA/DNA and DNA/RNA interaction since the reverse primer interacts with the SARS-CoV-2 genome along with the negative sense cDNA strand [3]

| Primer-probe | Mutation in ref. genome | Mutation Position | Primer Binding Energy (kcal/mol) | Interaction type |
| --- | --- | --- | --- | --- |
| CCDC-N-F | no mutation |  | -283.27 | DNA/DNA |
|  | G → A | 28,881 | -282.43 | DNA/DNA |
|  | G → A | 28,882 | -281.6 | DNA/DNA |
|  | G → C | 28,883 | -279.62 | DNA/DNA |
| CCDC-ORF1-F | no mutation |  | -138.05 | DNA/DNA |
|  | C → T | 13,358 | -133.77 | DNA/DNA |
| CCDC-ORF1-P | no mutation |  | -144.82 | DNA/DNA |
|  | T → G | 13,402 | -147.82 | DNA/DNA |
| E-Sarbeco-R | no mutation |  | -167.39 | RNA/DNA |
|  | no mutation |  | -167.77 | DNA/DNA |
|  | G → T | 26,370 | -149.57 | RNA/DNA |
|  | G → T | 26,370 | -149.49 | DNA/DNA |
| HKU-N-F |  |  | -279.36 | DNA/DNA |
|  | T → C | 29,148 | -275.41 | DNA/DNA |
| 2019-nCoV_N1-P | no mutation |  | -282.37 | DNA/DNA |
|  | C → T | 28,311 | -279.49 | DNA /DNA |
| 2019-nCoV_N1-R | no mutation |  | -277.67 | RNA/DNA |
|  | no mutation |  | -277.85 | DNA/DNA |
|  | C → A | 28,344 | -273.22 | RNA/DNA |
|  | C → A | 28,344 | -273.4 | DNA/DNA |
| 2019-nCoV_N3-F | no mutation |  | -280.31 | DNA/DNA |
|  | T → C | 28,688 | -279.86 | DNA/DNA |
| 2019-nCoV_N3-R | no mutation |  | -272.3 | RNA/DNA |
|  | no mutation |  | -273.04 | DNA/DNA |
|  | C → T | 28,739 | -267.95 | RNA/DNA |
|  | C → T | 28,739 | -268.57 | DNA/DNA |

#### Mutations that cause no change to the interaction structure

The mutations in the primer binding region of the CCDC-N-F at base pair positions 28,881 and 28,882 had no affect on the primer binding ability.

The mutation in the primer binding region of 2019-nCoV_N3-F mutation at base pair position 28,688 did not change the structure. The primer is still able to bind to the mutated base, but the final base at the 3’ end of the primer remains unpaired. However, this unpaired base is the same in the structure of the 2019-nCoV_N3-F primer interacting with the reference genome NC_045512.2.

#### Mutations that cause partial mismatching

The following mutations resulted in a single mismatch between the primer/probe and the mutated base: CCDC-ORF1-P at base position 13,402, 2019-nCoV_N1-R at base position 28, 344 and CCDC-N-F at base position 28, 883. The other bases in the probe/primer were still able to pair with their target nucleotides.

The mutation in the primer binding region of CCDC-ORF1-F at base position 13, 358 affects the ability of the 5 bases at the 3’ end of the primer to bind. The remaining 16/21 bases of the forward primer are still able to bind to their target nucleotides.

The mutation in the probe binding region of 2019-nCoV_N1-P at base position 28, 311 is predicted to cause a mismatch of the first three bases at the 5’ end of the probe. The remaining 21 nucleotides of the probe still bind as expected.

#### Mutations that cause complete mismatching

The mutation in the primer binding region of E-Sarbeco-R at base pair position 26,370 is predicted to result in a complete mismatch between the reverse primer and the SARS-CoV-2 genome. The primer does not bind to its target. There is some partial binding 600 bp downstream of the target region and partial base pairing of the primer to itself. The mutation also causes the same complete mismatch between the primer and the positive sense cDNA strand.

The mutation in the primer binding region of HKU-N-F at base position 29, 148 is predicted to result in complete mismatch between the forward primer and the negative sense cDNA strand.

The mutation in the primer binding region of 2019-nCoV_N3-R mutation at base position 28, 739 is predicted to result in a complete mismatch between the reverse primer and the SARS-CoV-2 RNA genome. The primer does not bind to the target area. However, the reverse primer is able to bind to the target area on the positive sense cDNA, except with one base pair mismatch at the mutated base.

## Discussion

We used DinoKnot to predict the secondary structure of the interaction between RNA transcripts from the SARS-CoV-2 genome and the reverse primers from nine primer-probe sets in order to see if the RNA structure of SARS-CoV-2 affected reverse primer binding. In addition, We predicted how mutations listed by Vogels *et al*. in the primer/probe binding regions may affect the binding ability of the primer/probes, and thus, SARS-CoV-2 detection. We found that all reverse primers were able to interact with their target regions and predicted partial mismatching to occur between some reverse primers. We also predicted three mutations to prevent the ability of the primer/probe to bind to their target region.

### Reverse primer interaction with the SARS-CoV-2 RNA genome

The following primers were predicted to have partial mismatching in the primer binding region: 2019-nCoV_N2-R, HKU-N-R, E-Sarbeco-R, and RdRp-SARSr-R. The 2019-nCoV_N2, E-Sarbeco and HKU-N primer-probe sets had analytical efficiencies and sensitivities that are similar to the other primer-probe sets [4]. The RdRp-SARSr primer-probe set had an analytical efficiency comparable to the other primer-probe sets but a reduced sensitivity [4].

The single base mismatch in the 2019-nCoV_N2-R primer is unlikely to affect the ability of the reverse transcriptase to generate the cDNA strand. The reverse transcriptase generates the cDNA strand in the 5’ to 3’ direction so a mismatch at the 3’ end is unlikely to affect strand synthesis. As well, this primer-probe sets had high analytical efficiency and sensitivity shown by Vogels *et al*. which supports that this mismatch is unlikely to have an effect on cDNA strand synthesis [4]

The mismatching in the E-Sarbeco-R and HKU-N-R primers occur when the primer is assumed to have structure prior to interaction with the target. However, when the primer was forced in DinoKnot as unfolded, the program predicts the E-Sarbeco-R primer to fully bind to its target sequence and the HKU-N -R primer to bind with a single mismatch at the 5’ end of the primer. Since the qRT-PCR assay used by Vogels *et al*. has a denaturation step of 95° C and an annealing step of 55° C, it can be assumed that the primer is fully unfolded at those temperatures because the E-Sarbeco-R and HKU-N-R had primer efficiencies comparable to the other primers as determined by Vogels *et al*. [4].

The RdRp-SARSr-R primer contains two degenerate bases, R and S. Vogels *et al*. determined that 990/992 of their SARS-CoV-2 clinical samples contain a T at the position in the reference genome NC_045512.2 where the S base binds to its target [4]. This primer-probe set had the lowest sensitivity out of all of the primers [4]. During the qRT-PCR test by Vogels *et al*. this primer-probe set had the highest Ct values, 6-10 Ct values higher than the other sets, and is unable to detect low viral amounts [4]. DinoKnot was run with all possible combinations of the position of the degenerate base R input as an A or G and S input as a C or G shown in Fig 3.

The combination that produced the expected match was when an A was input at the R position and a G was input at the S postion. Vogels *et al*. proposed that changing the S to an A in the reverse primer could increase the sensitivity of the primer-probe set [4]. To test this, the primer was given as input to DinoKnot with an A input at the S position. All combinations where S was input as an A resulted in the primer not binding to the target area. The result was the primer binding to itself and partial binding to the 5’ end of the transcript, over 400 bases downstream from the target area. Therefore, this result indicates that changing the S to an A is unlikely to increase the sensitivity of the primer even though the target base pair is a U in the RNA genome and a T on the cDNA transcript. Based on these results, we hypothesize that the low sensitivity issue of the RdRp-SARSr primer-probe set may be due to the predicted primer mismatch when a G is present in the R and S position of the primer. Although the concentration of each base combination at the degenerate R and S positions is not stated in the protocol, if the assumption is made that the four combinations are present in equal amounts, this may explain the low sensitivity issue of the RdRp primer probe-set [30] [4]. The base combination where a G is present at both the R and S position is predicted to mismatch to the target region. This would lower the RdRp-SARSr-R concentration that is capable of binding to its target region. If the effective concentration of the RdRp-SARSr-R primer is lower than the other primer-probe sets, this may explain why this set had low sensitivity issues due to higher Ct values shown experimentally by Vogels *et al*. [4]. Therefore, we suggest that changing the base at the R position to an A and the base at the S position to a G may increase the RdRp-SARSr primer-probe set sensitivity since this base combination was predicted to perfectly bind to its target region.

### Mutations in the expected primer/probe binding region

Vogels *et al*. looked at 992 clinical samples and identified mutations in the expected primer binding regions that could decrease the primer sensitivity for SARS-CoV-2 detection [4]. We used DinoKnot to predict how those mutations affect the ability of the primer/probe to bind to its target sequence. Mutations in the expected binding region of the forward primer and probe were DNA/DNA interactions and mutations in the reverse primer expected binding region show both the RNA/DNA interaction and the DNA/DNA interaction.

Based on the interaction structures predicted by DinoKnot, the mutations in the primer binding regions of the E-Sarbeco-R, N-HKU-F, and 2019-nCoV_N3-R primers are most likely to have the greatest effect on decreasing primer sensitivity for SARS-CoV-2 detection. This is because the interaction structures predicted by DinoKnot show no binding of the primer to the target area. All mutations resulted in an increase in the energy of binding, except for the mutation in the CCDC-ORF1-P binding region, which lowered the energy of binding. The energy increase means that binding is not as favourable as before and in environments where there is competition for binding, binding may not happen. The mutations in E-Sarbeco-R, N-HKU-F, and 2019-nCoV_N3-R binding regions caused the greatest increase in binding energy.

The mutation in the primer binding region of the E-Sarbeco-R primer at base position 26, 370 is predicted by DinoKnot to cause a complete mismatch between the reverse primer and the RNA genome, as well as between the reverse primer and positive sense cDNA strand. Therefore, this single base mutation may cause the mutated SARS-CoV-2 strains to not be detected by the E-Sarbeco primer-probe set, especially since the reverse primer is needed for the generation of the cDNA strand. Even if the forward primer and probe can still bind as expected to their target sequence, if the reverse primer cannot bind to the RNA genome, the reverse transcriptase will not be able to generate the negative sense cDNA strand for the forward primer to bind to.

The mutation in the HKU-N-F primer binding region at position 29, 148 was predicted by DinoKnot to result in the forward primer not being able to bind to its target region. Without a functional forward primer, this would result in the inability to generate a negative sense cDNA copy, and therefore no amplification of either the positive sense or negative sense cDNA strand. As shown in Fig 1, the negative sense cDNA is required for the generation of the positive sense cDNA strand and any subsequent amplification. Even if the reverse primer and probe bind as expected to their target region, there would only be the generation of one positive sense cDNA strand and no amplification.

The mutation in the 2019-nCoV_N3-R primer binding region at position 28, 739 is predicted to cause a complete mismatch between the reverse primer and the RNA genome. However, the reverse primer is able to bind to the target area on the positive sense cDNA, except with one base pair mismatch at the mutated base. This mutation may cause this strain not to be detected by this primer-probe set because if the reverse primer does not bind to the target area on the SARS-CoV-2 genome, the reverse transcriptase cannot generate the positive sense cDNA strand. Therefore, the binding of the reverse primer to the negative sense cDNA strand interaction would not occur.

The CCDC-ORF1-F primer had a mismatch at the 3’ end of the primer causing the 5 bases at 3’ end to remain unpaired. The DNA polymerase generates the complementary strand in the 5’-3’ direction. Since the 5’ end still binds, the full target area may still be generated. In this primer-probe set, the probe binds to the same area of the cDNA that the forward primer binds so having the full area of the transcript generated is important. The 5 bases at the 3’ end did not align to any other part of the reference genome in a NCBI BLAST search so it is unlikely but still possible that the tail could bind to another part of the genome and hinder the ability of the DNA polymerase to generate the cDNA strand, preventing SARS-CoV-2 detection [29]. This mutation should be tested experimentally in order to determine its effect on the primer-probe set’s ability for SARS-CoV-2 detection.

### Viral Load and SARS-CoV-2 detection

Recent work by Pan *et al*. on 82 infected individuals with SARS-CoV-2 determined the viral load in sputum and swab samples to peak on day 5-6 after symptom onset, with a range of 10^4^ to 10^7^ copies per mL [34]. This is earlier than SARS-CoV which peaks approximately 10 days after symptom onset [34]. Vogels *et al*. determined the qRT-PCR primer-probe sets to have a limit of detection of 10^5^ copies per mL except for the RdRp-SARSr primer-probe set which tested negative in samples containing 10^0^-10^2^ viral RNA copies/μL concentrations (which is equivalent to 1-100 viral RNA copies/mL) [4]. Pan *et al*. determined that early after the onset of symptoms, the viral load is greater than 1 × 10^6^ copies per mL [34]. This is sufficient for detection by the primer-probe sets, except for RdRp-SARSr [4]. However, respiratory samples from 80 patients from different stages of infection showed a range of 641 copies per mL to 1.34 × 10^11^ copies per mL. Therefore, the more time that passes after symptom onset may result in the viral load to be below the limit of detection during testing. If patients are tested a greater number of days after symptom onset when the viral load may be below the primer-probe set limit of detection determined by Vogels *et al*., this could result in a false negative test. However, nasopharyngeal swabs are the most common ways to collect patient samples and the work by Pan *et al*. only looked at one nasal swab which was tested 3 days after symptom onset with a total of 1.69 × 10^5^ copies per mL [2, 34].

In a patient led survey, 27.5% of respondents believed they had COVID-19 but tested negative for the disease [35]. This group of respondents were on average tested later, by day 16, compared to the 21% of respondents who believed they had COVID-19 and also tested positive, who were on average tested by day 10 [35]. As is a limitation of this survey we cannot verify whether the respondents who believed they had COVID-19 were in fact infected with SARS-CoV-2. However, since they were tested later, this may support the hypothesis that the viral load may be below the limit of detection if the test is taken a greater number of days after symptom onset. Therefore false negative tests are likely an issue with having enough viral load in the sample rather than an issue with the primer interaction with the SARS-CoV-2 genome since DinoKnot predicted correct primer binding for most of the primers at the higher temperatures.

We hypothesize that a SARS-CoV-2 aptamer test is beneficial to provide a lower limit of detection. Aptamers are single stranded RNA or DNA nucleotides (10-100nt) that are able to bind to targets such as viruses and proteins [36]. Aptamers’ binding specificity is ensured by their secondary and tertiary structure [36]. An RNA aptamer test designed for the SARS-CoV Nucleocapsid protein showed a detection limit of 2 pg/mL [37]. A recent aptamer test for Norovirus, a positive sense RNA virus, has a limit of detection of 200 viral copies/mL [38]. This is lower than the qRT-PCR limit of detection of determined by Vogels *et al*. by a magnitude of 10^3^ [4]. The lower limit of detection that is possible with aptamer tests would be beneficial in testing patients later after symptom onset.

We believe that an aptamer test can be designed for SARS-CoV-2 that both improves the detection sensitivity and provides a more rapid test compared to the qRT-PCR tests currently in use. The areas targeted in the primer-probe sets may be good targets for aptamer design, since they are specific to SARS-CoV-2. Zooming in on these regions in the GISAID h-CoV-19 app, it can be shown that these areas generally have few mutations, as compared to the number of mutations that can be seen in the entire genome as shown in Fig 5 [33].

**Fig 5.**
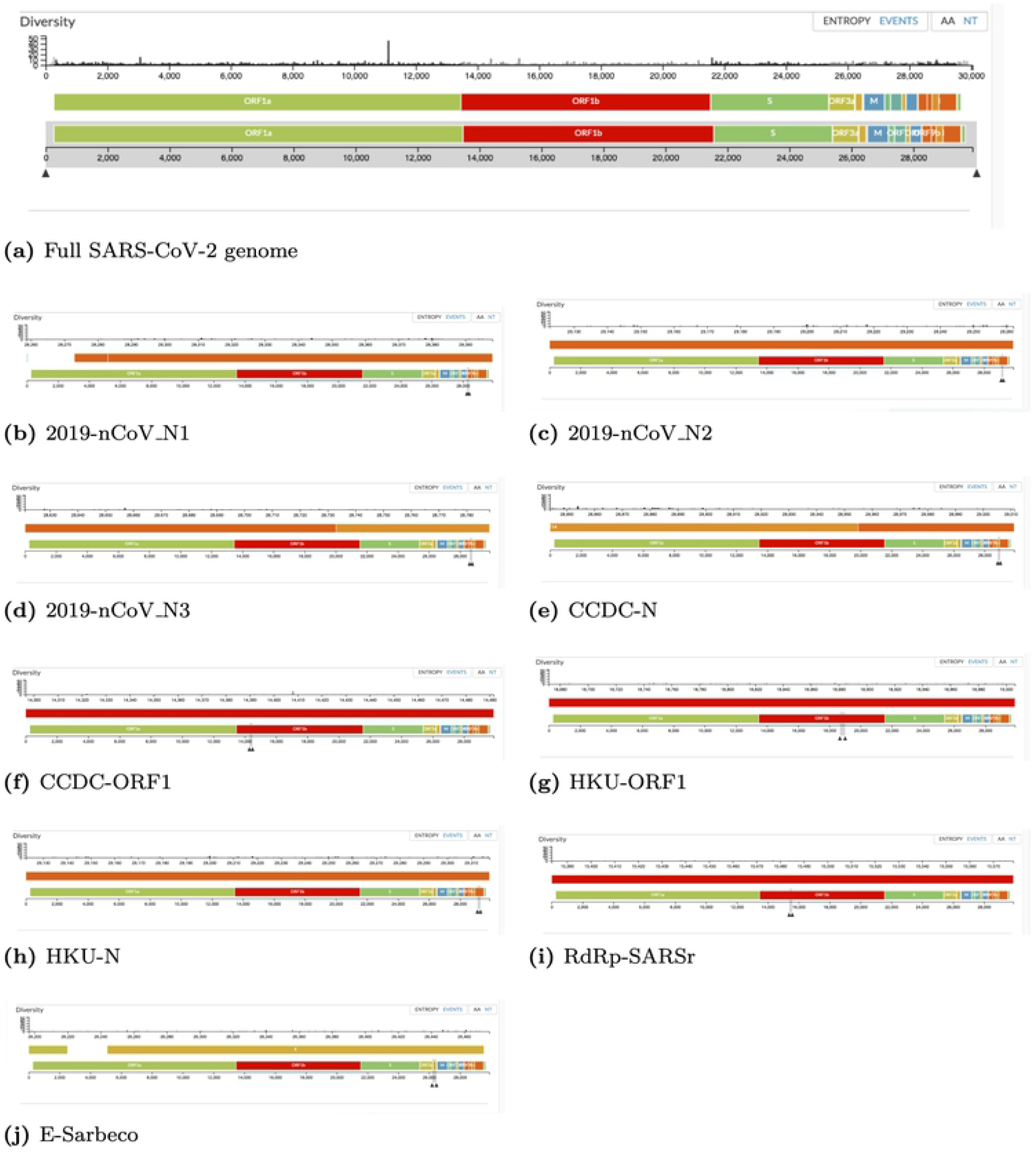
Screenshots from GISAID hCoV-19 genetic epidemiology mutation events. Mutation events reported to GISAID for nucleic acid positions of the (a) full SARS-CoV-2 genome and (b-j) zoomed in sections of the primer-probe sets as of July 23, 2020.

Our findings may be beneficial in a starting place for aptamer design. The HKU-ORF1 primer-probe set was the only set to have its reverse primer completely bind its target sequence and not contain any mutations in the primer/probe target areas in the 992 clinical samples looked at by Vogels *et al*. [4]. The GISAID h-CoV-19 app also shows the area targeted by this primer-probe set to have a low number of mutation events [33]. The 2019-nCoV_N2 primer-probe set did not have any mutations in the primer/probe binding regions listed by Vogels *et al*. but the reverse primer did contain the one base pair mismatch to the reference genome. The areas targeted by these primer-probe sets may be suitable targets for designing a SARS-CoV-2 aptamer test since these areas are specific for SARS-CoV-2 and contain few mutations.

## Conclusion

We introduced the program DinoKnot that is able to predict the secondary structure of two interacting nucleotide sequences. By in silico prediction we provided evidence to support that during COVID-19 qRT-PCR tests, the reverse primers investigated here are binding to their expected target region on the SARS-CoV-2 genome. Our prediction supports that binding is not prevented by the secondary structure of the SARS-CoV-2 RNA genome. This supports the idea that the correct area of the SARS-CoV-2 genome is being amplified during the qRT-PCR tests and the RNA structure does not prevent SARS-CoV-2 detection. Our prediction also provides a hypothesis to explain the low sensitivity issue of the RdRp-SARSr primer-probe set and we suggest changing the degenerate base positions in the RdRp-SARSr-R from an R to an A and an S to a G to potentially increase the sensitivity.

We also predicted how different mutations in the primer binding region affected primer binding and that the SARS-CoV-2 strains with mutations in the E-Sarbeco-R, N-HKU-F, and 2019-nCoV_N3-R primer binding regions may reduce the sensitivity of these COVID-19 tests for SARS-CoV-2 detection.

Finally we hypothesize that an aptamer test may be a better way to test for COVID-19 since it can provide a lower limit of detection. Further research is required to quantify the viral load at each stage of infection and determine at what point the qRT-PCR can no longer detect COVID-19 so that conclusions can be made on the impact of the limit of detection affecting false negative test results.

DinoKnot can also be used as a tool to aid in the prevention of false positive results during the design COVID-19 and other nucleic acid based testing. A false positive test would involve the non-specific binding of the primer-probe sets to a source other than the target in the patient sample. Future work can be done using our program DinoKnot to predict the interaction between primer/probes with other pathogens and nucleic acid sequences that may be present in patient samples to determine what the primer/probes may be capable of binding to produce false positive results.

## References

1. Zhou P, Yang X, Wang X, et al. A pneumonia outbreak associated with a new coronavirus of probable bat origin. Nature. 2020;579:270–273. doi:10.1038/s41586-020-2012-7.

2. Wang W, Xu Y, Gao R, et al. Detection of SARS-CoV-2 in Different Types of Clinical Specimens. JAMA. 2020;323:1843–1844. doi:10.1001/jama.2020.3786.

3. Bustin SA, Mueller R. Real-time reverse transcription PCR (qRT-PCR) and its potential use in clinical diagnosis. Clin Sci (Lond). 2020;109:365–379. doi:10.1042/CS20050086.

4. Vogels CBF, Brito AF, Wyllie AL, et al. Analytical sensitivity and efficiency comparisons of SARS-CoV-2 RT–qPCR primer–probe sets. Nat Microbiol. 2020;doi:10.1038/s41564-020-0761-6.

5. Andronescu M, Aguirre-Hernández R, Condon A, Hoos HH. RNAsoft: A suite of RNA secondary structure prediction and design software tools. Nucleic Acids Res. 2003;31(13):3416–3422. doi:10.1093/nar/gkg612.

6. Andronescu M, Chuan Z, Condon A. Secondary structure prediction of interacting RNA molecules. Journal of molecular biology. 2005;345(5):987–1001. doi:10.1016/j.jmb.2004.10.082.

7. Dirks RM, Bois JS, Schaeffer JM, Winfree E, Pierce NA. Thermodynamic Analysis of Interacting Nucleic Acid Strands. SIAM Rev. 2007;49(1):65–88. doi:10.1137/060651100.

8. Alkan C, Karakookay E, Nadeau JH, Sahinalp SC, Zhang K. RNA-RNA Interaction Prediction and Antisense RNA Target Search. Journal of Computational Biology. 2006;13(2):267–282. doi:10.1089/cmb.2006.13.267.

9. Kato Y, Akutsu T, Seki H. A grammatical approach to RNA-RNA interaction prediction. Pattern Recognition. 2009;42(4):531–538. doi:10.1016/j.patcog.2008.08.004.

10. Chitsaz H, Salari R, Sahinalp SC, Backofen R. A partition function algorithm for interacting nucleic acid strands. Bioinformatics. 2009;25(12):i365–i373. doi:10.1093/bioinformatics/btp212.

11. Dimitrov RA, Zuker M. Prediction of Hybridization and Melting for Double-Stranded Nucleic Acids. Biophysical Journal. 2004;87(1):215–226. doi:10.1529/biophysj.103.020743.

12. Rehmsmeier M, Steffen P, Hochsmann M, Giegerich R. Fast and effective prediction of microRNA/target duplexes. RNA (New York, NY). 2004;10(10):1507–1517. doi:10.1261/rna.5248604.

13. Lorenz R, Bernhart SH, Honer zu Siederdissen C, Tafer H, Flamm C, Stadler PF, et al. ViennaRNA Package 2.0. Algorithms for Molecular Biology. 2011;6(1):26. doi:10.1186/1748-7188-6-26.

14. Reuter JS, Mathews DH. RNAstructure: software for RNA secondary structure prediction and analysis. BMC bioinformatics. 2010;11(1):129+. doi:10.1186/1471-2105-11-129.

15. Kery MB, Feldman M, Livny J, Tjaden B. TargetRNA2: identifying targets of small regulatory RNAs in bacteria. Nucleic acids research. 2014;42(Web Server issue):W124–W129. doi:10.1093/nar/gku317.

16. Nussinov R, Jacobson AB. Fast algorithm for predicting the secondary structure of single-stranded RNA. Proceedings of the National Academy of Sciences of the United States of America. 1980;77(11):6309–6313. doi:10.1073/pnas.77.11.6309.

17. Zuker M, Stiegler P. Optimal computer folding of large RNA sequences using thermodynamics and auxiliary information. Nucleic acids research. 1981;9(1):133–148. doi:10.1093/nar/9.1.133.

18. Akutsu T. Dynamic programming algorithms for RNA secondary structure prediction with pseudoknots,. Discrete Applied Mathematics. 2000;104(1-3):45–62. doi:10.1016/s0166-218x(00)00186-4.

19. Lyngsø RB, Pedersen CNS. Pseudoknots in RNA secondary structures. In: Proceedings of the fourth annual international conference on Computational molecular biology. RECOMB ‘00. New York, NY, USA: ACM; 2000. p. 201–209. Available from: http://dx.doi.org/10.1145/332306.332551.

20. Sheikh S, Backofen R, Ponty Y. Impact of the Energy Model on the Complexity of RNA Folding with Pseudoknots. In: Kärkkäinen J, Stoye J, editors. Combinatorial Pattern Matching. vol. 7354 of Lecture Notes in Computer Science. Springer Berlin Heidelberg; 2012. p. 321–333. Available from: http://dx.doi.org/10.1007/978-3-642-31265-6_26.

21. Rivas E, Eddy SR. A dynamic programming algorithm for RNA structure prediction including pseudoknots. Journal of molecular biology. 1999;285(5):2053–2068. doi:10.1006/jmbi.1998.2436.

22. Dirks RM, Pierce NA. A partition function algorithm for nucleic acid secondary structure including pseudoknots. J Comput Chem. 2003;24(13):1664–1677. doi:10.1002/jcc.10296.

23. Reeder J, Giegerich R. Design, implementation and evaluation of a practical pseudoknot folding algorithm based on thermodynamics. BMC Bioinformatics. 2004;5(1):104+. doi:10.1186/1471-2105-5-104.

24. Jabbari H, Condon A, Zhao S. Novel and efficient RNA secondary structure prediction using hierarchical folding. J Comput Biol. 2008;15(2):139–163. doi:10.1089/cmb.2007.0198.

25. Jabbari H, Wark I, Montemagno C, Will S. Knotty: efficient and accurate prediction of complex RNA pseudoknot structures. Bioinformatics. 2018;34(22):3849–3856. doi:10.1093/bioinformatics/bty420.

26. Jabbari H, Condon A. A fast and robust iterative algorithm for prediction of RNA pseudoknotted secondary structures. BMC Bioinformatics. 2014;15(1):147+. doi:10.1186/1471-2105-15-147.

27. Tinoco I, Bustamante C. How RNA folds. Journal of Molecular Biology. 1999;293(2):271–281. doi:10.1006/jmbi.1999.3001.

28. Andronescu M, Condon A, Hoos HH, Mathews DH, Murphy KP. Computational approaches for RNA energy parameter estimation. RNA. 2010;16(12):2304–2318. doi:10.1261/rna.1950510.

29. Coordinators NR. Database resources of the National Center for Biotechnology Information. Nucleic Acids Res. 2016;44:(D1):D7–D19. doi:10.1093/nar/gkv1290.

30. WHO. WHO - Coronavirus disease (COVID-19) technical guidance: Laboratory testing for 2019-nCoV in humans.; 2020 (Retrieved Sept 6, 2020). Available from: https://github.com/HosnaJabbari/DinoKnot_data/blob/master/whoinhouseassays.pdf.

31. Iserte JA, Stephan BI, Goñi SE, Borio CS, Ghiringhelli PD, Lozano ME. Family-Specific Degenerate Primer Design: A Tool to Design Consensus Degenerated Oligonucleotides. Biotechnology Research International. 2013;2013:9. doi:10.1155/2013/383646.

32. Darty K, Denise A, Ponty Y. VARNA: Interactive drawing and editing of the RNA secondary structure. Bioinformatics. 2009; p. 1974–1975.

33. Elbe S, Buckland-Merrett G. Data, disease and diplomacy: GISAID’s innovative contribution to global health.; 2017.

34. Pan Y, Zhang D, Yang P, Poon LLM, Wang Q. Viral load of SARS-CoV-2 in clinical samples. The Lancet - Correspondence. 2020;20:411–412. doi:10.1016/S1473-3099(20)30113-4.

35. Assaf G, Davis H, McCorkell L, Wei H, Brooke O, Akrami A, et al. An Analysis of the Prolonged COVID-19 Symptoms Survey by Patient-Led Research Team; 2020. Available from: https://patientresearchcovid19.com/research/report-1/.

36. Zou X, Wu J, Gu J, Shen L, Mao L. Application of Aptamers in Virus Detection and Antiviral Therapy. Front Microbiol. 2019;10:1462. doi:10.3389/fmicb.2019.01462.

37. Ahn DG, Jeon IJ, Kim JD, Song MS, Han SR, Lee SW, et al. RNA aptamer-based sensitive detection of SARS coronavirus nucleocapsid protein. Analyst. 2009;134:1896–1901. doi:10.1039/B906788D.

38. Weerathunge P, Ramanathan R, Torok VA, Hodgson K, Xu Y, Goodacre R, et al. Ultrasensitive Colorimetric Detection of Murine Norovirus Using NanoZyme Aptasensor. Anal Chem. 2019;91:3270–3276.

